# Accurate and scalable decontamination of imaging-based spatial transcriptomics via optimal transport

**DOI:** 10.64898/2026.09.09.750350

**Authors:** Yuheng Chen, Yuyao Liu, Zitong Chao, Shi Han, Yeqin Zeng, Baichen Yu, Fan Zhang, Angela Ruohao Wu, Jiguang Wang, Hao Chen, Jiashun Xiao, Can Yang

## Abstract

Imaging-based spatial transcriptomics enables molecule-resolved profiling of gene expression and tissue organization in situ. However, segmentation errors, transcript spillover and three-dimensional cell overlap can introduce misassigned transcripts into cell-level expression profiles, compromising biological interpretation and obscuring genuine signals. Existing methods either remove suspect expression at the cost of signal loss or lack a biologically grounded criterion for transcript assignment. Here we present CellDot, an optimal-transport framework that determines the fate of each transcript by retaining it in its host cell, reassigning it to a plausible neighboring cell or removing it as background. By integrating reference-guided expression compatibility with spatial information and data-adaptive constraints, CellDot enables accurate and traceable molecule-level correction while preserving biologically meaningful variation. In evaluations across multiple human tumor datasets, CellDot exhibited superior performance compared to existing decontamination methods, successfully restoring spatial expression patterns that matched independent cross-platform measurements. Moreover, it significantly enhanced the recovery of cellular states, intercellular communication, and spatial niche programs. Our experiments using real data demonstrated CellDot’s scalability and established it as the only method applicable to a whole-transcriptome Atera dataset, underscoring its distinct advantages in the field of spatial transcriptomics.

## Introduction

A central question in biology is how cells are spatially organized and how they interact with one another[1]. Spatial transcriptomics (ST) enables the measurement of gene expression within intact tissues, facilitating the construction of comprehensive maps of cell types, states, spatial distributions, and interactions. Among ST technologies, imaging-based approaches[2–5], such as Xenium, represent an important class that directly resolves individual RNA molecules with their molecular identities and spatial coordinates. This molecule-resolved capability provides an unprecedented opportunity to investigate tissue organization at cellular and subcellular resolution. Recently, the release of the Atera platform by 10x Genomics has further extended imaging-based ST toward whole-transcriptome profiling while maintaining molecule-level spatial resolution[6], opening new possibilities for comprehensive characterization of cellular states, spatial organization, and molecular interactions across tissues.

Despite these advantages, transcript contamination remains a major limitation for accurately interpreting cell-level expression profiles, where RNA molecules originating from one cell are incorrectly assigned to and counted as part of another cell’s genuine expression profile. In imaging-based ST analysis, detected transcripts are typically assigned to cells based on segmented cell boundaries. However, perfect cell segmentation may not be achievable due to factors such as weak membrane staining, irregular cell morphology, and the densely packed nature of cells. As a result, RNA molecules located near cell boundaries can be mistakenly assigned to neighboring cells[7–9]. In addition, transcripts may diffuse away from their original locations during tissue processing, causing transcripts to enter the boundaries of other cells or accumulate in extracellular regions without a clear cellular origin. More fundamentally, tissues are inherently three dimensional, so the intricate arrangement of cells across the depth of a tissue section makes overlap inevitable, causing transcripts from one cell to fall within another cell’s projected two-dimensional boundary and obscuring their true cellular origin[10]. Together, these sources of contamination introduce false-positive expression signals into cell-level profiles that can propagate into downstream analyses, leading to spurious biological interpretations such as the identification of cell states defined by neighboring cell markers, assignment of cell–cell communication signals to incorrect sender cells, and artificial spatial programs driven by adjacent cells’ transcripts. The accumulation of such erroneous signals may further obscure subtle but genuine biological signals, reducing the ability to detect biologically meaningful differences and spatial patterns[11, 12].

Although several methods have been developed to address transcript contamination, major limitations still restrict their effectiveness and practical utility. First, some methods operate directly on the assembled cell-level expression profile and therefore discard transcript-level information, preventing misplaced or unassigned transcripts from being traced back and reassigned to their true cellular origins[11, 13, 14]. Such approaches may therefore overcorrect the data by removing genuine biological signal together with contamination and sacrifice molecule-level traceability. Second, existing transcript-level methods often lack a biologically grounded criterion for determining the true cellular origin of each molecule[12, 15]. Instead, transcript fates are inferred indirectly from objectives such as improving cell type separability or identifying expression patterns that appear foreign to the host cell, which may not provide sufficient guidance for accurate reassignment and can leave residual contamination. A further challenge is scalability, as imaging-based ST is rapidly expanding from targeted panels to whole-transcriptome profiling (e.g., the Atera platform[6]), with individual tissue sections containing millions of cells and billions of molecules.

To overcome these limitations and provide an accurate and scalable solution for transcript-level decontamination, we present CellDot (**Cell D**econtamination by **O**ptimal **T**ransport). CellDot formulates transcript correction as a capacitated entropic optimal transport problem[16–19], where each transcript can either remain in its segmented host cell, be reassigned to a nearby cell with compatible gene expression, or be discarded as ambient background noise when no plausible cellular origin can be identified. Guided by a paired single-cell reference that provides a reliable biological prior for transcript ownership, CellDot improves the accuracy and robustness of the decontamination process. More importantly, beyond transcript-level preferences, CellDot incorporates capacity constraints to prevent excessive reassignment or removal, ensuring biologically plausible transcript assignment. This formulation leads to a convex optimization problem with a unique optimal solution, enabling efficient scaling to whole-transcriptome datasets while providing an accurate and traceable fate assignment for every molecule in the raw data.

We demonstrate CellDot across human tumor sections spanning multiple cancer types and diverse gene panel sizes, ranging from targeted panels to whole-transcriptome measurements. CellDot achieves superior overall performance in transcript correction and cell-type characterization compared with existing approaches[11–15], while scaling to the large whole-transcriptome Atera dataset[6] which cannot be handled by existing methods. In colorectal cancer, cross-platform validation against matched Visium HD[20] and Visium[21] measurements demonstrates that CellDot restores true spatial expression patterns and improves the accuracy of downstream gene imputation. CellDot also removes macrophage subpopulations driven by neighboring cell contamination, recovering biologically meaningful macrophage states. In lung adenocarcinoma, CellDot restores the correct cellular origin of ligand transcripts, removing spurious sender assignments and recovering biologically supported cell–cell communication patterns obscured by contamination. In cervical cancer profiled by the Atera platform, CellDot extends correction beyond targeted panels to whole-transcriptome measurements, enabling scalable removal of contamination-derived signals while preserving genuine lineage signals and restoring cell-intrinsic spatial programs. We envision CellDot as an essential decontamination step between segmentation[8, 22–25] and downstream analysis, restoring transcript ownership at the molecular level and enabling more reliable discovery of cellular states, interactions, and spatial organization.

## Results

### Overview of CellDot

CellDot performs decontamination on imaging-based ST data that have already undergone cell segmentation and annotation. It takes transcript identities and coordinates, cell boundaries, cell-type annotations, and a paired single-cell reference as input, and determines the final fate of each transcript. CellDot assumes that each detected transcript originates from one of three possible sources, namely its segmented host cell, a nearby cell, or the ambient background. Accordingly, each transcript is assigned one of three corresponding fates, remaining in its host cell, being reassigned to a nearby cell that better explains its presence, or being discarded when no plausible cellular origin can be identified (Fig. 1).

**Fig. 1.**
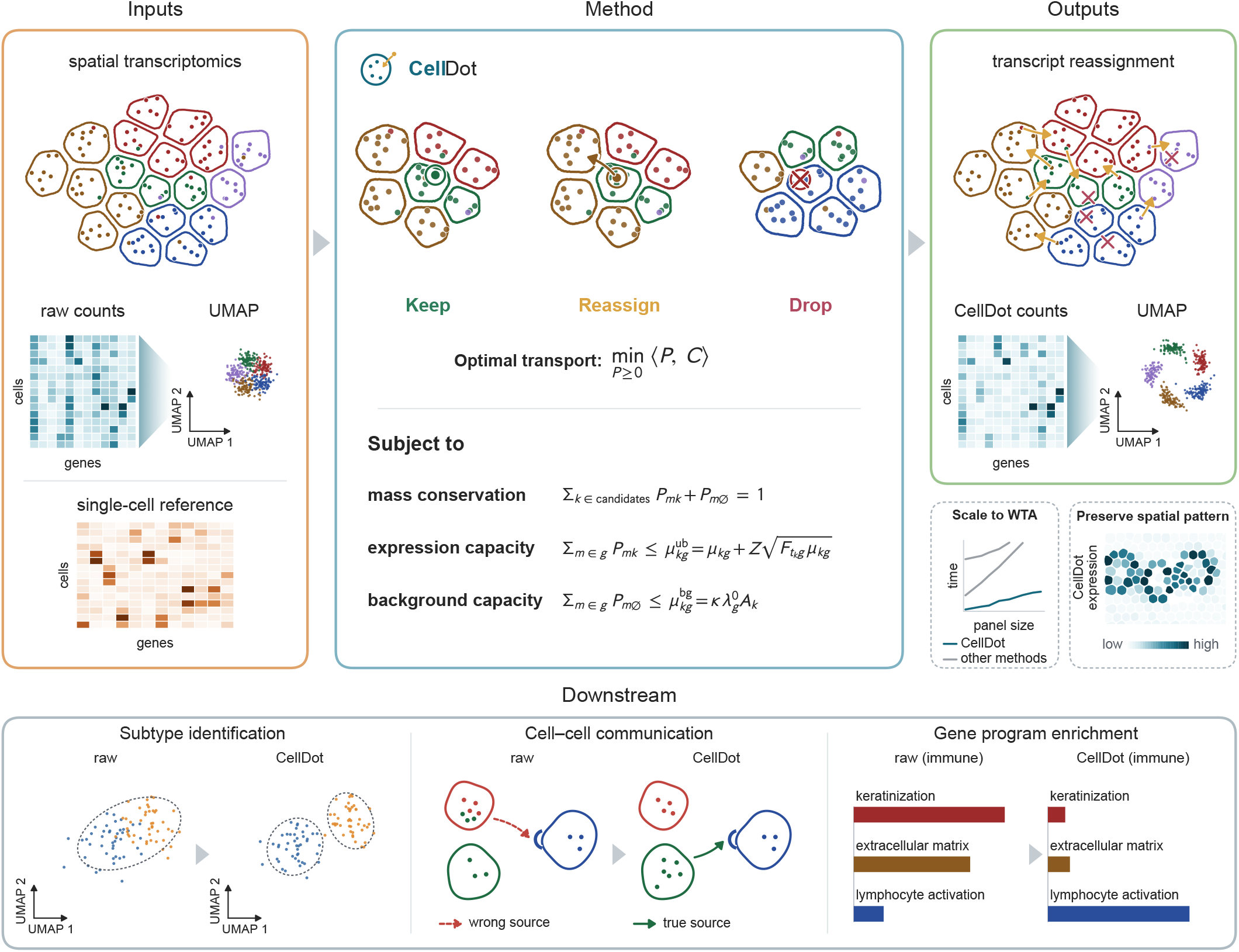
Overview of CellDot. Inputs, CellDot takes the coordinates and gene identities of all detected transcripts, the segmented cell boundaries and cell-type annotations, together with an annotated single-cell reference of the same tissue; the transcripts and boundaries define the contaminated cell × gene matrix. **Method**, each transcript is kept in its host cell, reassigned to a neighboring cell, or dropped as background. All fates are resolved jointly by an optimal-transport problem, in which a transcript is drawn to nearby cells whose type expresses its gene in the reference, and to the background when no nearby cell can explain it. Three constraints bound the solution: each transcript is placed exactly once (mass conservation), a cell can hold no more of a gene than its type is expected to express (expression capacity), and no more of a gene can be dropped than the measured background level allows (background capacity). **Outputs**, a traceable fate for every transcript and the corrected cell × gene matrix those fates define. Dashed insets, CellDot scales to whole-transcriptome-assayed (WTA) panels and preserves the spatial pattern of gene expression. **Downstream**, correction improves the analyses built on the corrected profiles: subtype identification recovers real populations rather than contamination artifacts, cell–cell communication is traced to its true senders, and gene programs are scored on a cell’s own transcripts.

CellDot formulates transcript assignment as an optimal transport problem, where transcripts are treated as sources and candidate cells in their local spatial neighborhoods, together with the ambient background, as possible sinks. The transport cost jointly considers spatial proximity and expression compatibility, with the latter guided by the single-cell reference. Importantly, CellDot calibrates this reference-derived prior to the spatial data using gene-specific scaling factors that account for platform-specific differences in gene detection efficiency[26] (Supplementary Fig. S1). The background sink is included to capture transcripts that cannot be plausibly explained by nearby cells, with its gene-specific background level estimated from transcripts detected in cell-free regions of the tissue section. This provides an empirical estimate of the background abundance of each gene and calibrates the likelihood of assigning a transcript to the background.

Moreover, to ensure biologically plausible transcript reassignment while preventing overcorrection, CellDot introduces two capacity constraints that are adaptively calibrated from the data (Fig. 1). The expression capacity is calibrated from the single-cell reference together with the observed transcript abundance of each cell, with a dispersion-aware margin that prevents excessive accumulation in recipient cells while preserving genuine high expression. The background capacity is estimated from gene-specific extracellular background density and the spatial area of each cell, bounding the amount of transcript mass that can be assigned to the background and thereby preventing excessive removal of measured transcripts (Supplementary Fig. S2).

CellDot solves the resulting convex optimal-transport problem using alternating Bregman projections[17, 18] in a tile-wise manner. This method ensures that computational costs remain linear with respect to the number of transcripts, while memory usage is optimized with respect to the size of individual tiles. As a result, CellDot achieves remarkable efficiency, enabling seamless scalability to entire tissue sections and large-scale whole-transcriptome datasets (Fig. 1). The molecule-level assignments generated by CellDot yield de-contaminated expression profiles that significantly enhance the quality of downstream analyses. These refined profiles enable more accurate subtype identification, robust inference of cell–cell communication, and deeper insights into gene programs, ultimately leading to a more faithful recovery of genuine biological signals (Fig. 1). CellDot is available as an open-source package at https://github.com/YangLabHKUST/CellDot, together with an interactive viewer of its molecule-level results on the four datasets analyzed in this paper at https://viewer.celldot.online/.

### CellDot outperforms baseline methods and scales to whole-transcriptome-assayed panels

To evaluate the performance of CellDot, we compared it with five existing decontamination methods, including resolVI[13], MisTIC[15], DenoIST[14], SPLIT[11] and cellAdmix[12], across multiple datasets from different tissue types spanning a broad range of panel sizes (Methods).

One major consequence of transcript contamination is the reduced separability between cell types, as transcripts originating from neighboring cell types can enter the same cellular expression profile and blur cell-type-specific expression patterns. On a Xenium breast cancer section (313-gene panel), the raw expression profiles yielded an average silhouette width (ASW) of only 0.034. While most methods increased the ASW, indicating improved separation of cell-type-specific expression patterns, CellDot achieved the largest improvement, raising the ASW to 0.224 and clearly outperforming the baseline methods (Fig. 2a,b). To provide a more comprehensive comparison, we further evaluated the mutually exclusive marker co-expression rate (MECR), with lower values indicating less cross-cell-type marker co-expression, and the positive marker purity (PMP), with higher values indicating greater concentration of cell-type markers within their corresponding cell types. CellDot achieved the lowest MECR and a PMP comparable to DenoIST, outperforming the other baselines and the raw data (Fig. 2b). However, DenoIST discarded approximately 27% of all measured transcripts on this dataset, substantially more than all other methods, suggesting that its high PMP may partly reflect more aggressive transcript removal rather than more accurate decontamination. Moreover, DenoIST showed a markedly lower ASW than CellDot. Similar results were observed on a Xenium colorectal cancer section (422-gene panel), where CellDot again achieved stronger overall performance, with a higher ASW and PMP and a lower MECR than all baseline methods (Supplementary Fig. S3).

**Fig. 2.**
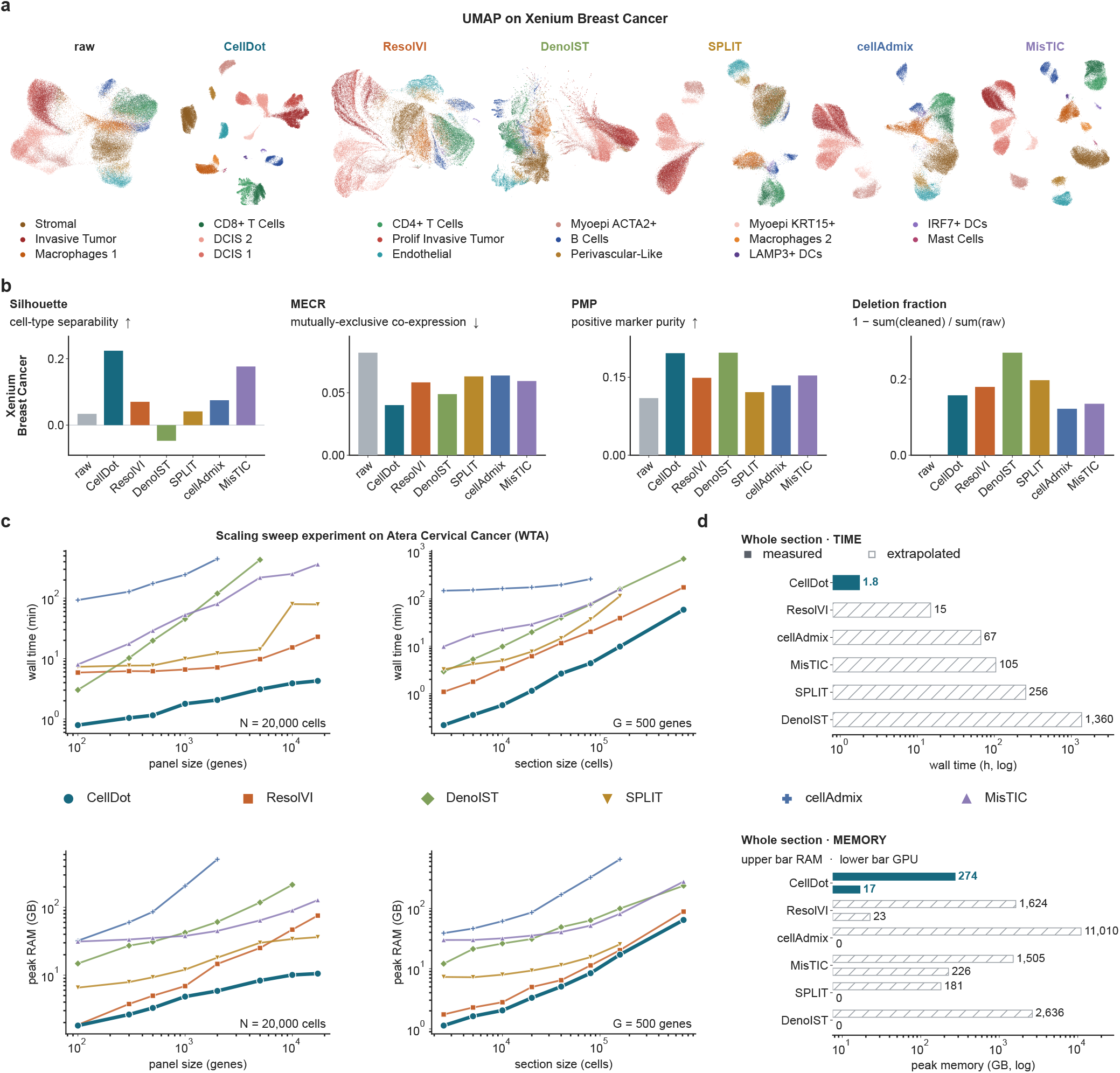
Benchmarking decontamination performance and scalability of CellDot against baseline methods. **a**, UMAPs of the Xenium breast cancer section (161,909 cells) computed on the raw counts and on the output of each method, all colored by the same cell-type annotation. **b**, four evaluation measures on this section: average silhouette width on the cell-type labels (higher is better), the mutually exclusive co-expression rate (MECR, lower is better), positive marker purity (PMP, higher is better) and the deletion fraction, defined as 1 − sum(corrected) / sum(raw). CellDot reaches the highest silhouette and the lowest MECR; its PMP is matched only by DenoIST, which deletes 27 % of the section’s transcripts against CellDot’s 16 % and returns the only negative average silhouette width. **c**, wall time (top) and peak RAM (bottom) against panel size at 20,000 cells (left) and against section size at 500 genes (right), on datasets subsampled from the whole-transcriptome cervical section. Both axes logarithmic; all runs measured on one host. **d**, run time (top) and peak memory (bottom) on the full cervical section, 717,576 cells × 17,285 genes. Solid bars, measured; hatched bars, extrapolated from the top of the same method’s own curve in **c**. In the memory panel the lower bar of each pair is peak GPU memory. CellDot is the only method that completes, correcting the section’s 880.8 million transcripts in 1.8 h with 274 GB of host RAM and 16.8 GB of GPU memory.

We next extended the comparison to a Xenium 5K lung adenocarcinoma dataset (5,001-gene panel). DenoIST and cellAdmix failed to complete due to computational constraints. CellDot again achieved the highest ASW among the remaining methods (Supplementary Fig. S3). Notably, resolVI achieved a lower MECR and a higher PMP than CellDot on this dataset. However, this performance was accompanied by the removal of 54% of all transcripts, compared with only 6% for CellDot, suggesting substantial overcorrection (Supplementary Fig. S3). Consequently, the median number of detected genes per cell dropped from 198 to 30 after resolVI correction, whereas CellDot retained 182 genes per cell (Supplementary Fig. S4).

To distinguish effective decontamination from simple transcript depletion, we further examined how correction reshaped the expression and co-expression patterns of cell-type markers. Across all three sections, CellDot consistently concentrated marker expression within the corresponding cell types while reducing ectopic expression in other populations (Supplementary Figs. S5–S7). In contrast, residual contamination remained evident after most competing methods, with markers of populations such as stromal cells still broadly detected in unrelated cell types. This pattern was further supported by differential expression analysis between the raw and corrected profiles, in which the genes showing the largest reductions within a given cell type were predominantly markers of other cell types (Supplementary Fig. S8). We next assessed marker co-expression using CS-CORE[27] in the colorectal cancer section. CellDot preserved co-expression among markers associated with the same cell type while reducing spurious co-expression across different cell types. In contrast, DenoIST weakened within-type co-expression, consistent with excessive removal of genuine signal, whereas resolVI introduced additional cross-type co-expression (Supplementary Fig. S9).

As imaging-based ST technologies continue to expand in gene panel size, computational scalability in both memory and runtime has become increasingly important. To evaluate the scalability of different methods, we used a whole-transcriptome cervical cancer dataset generated with the recently introduced Atera platform. CellDot showed approximately linear growth in runtime and memory with the number of cells and at most linear growth with the number of genes, maintaining the feasibility of whole-transcriptome correction (Fig. 2c). Across all tested panel sizes and section sizes, CellDot was consistently the fastest and most memory-efficient method, running three to six times faster than the closest competing approach. More importantly, on the full section, comprising 717,576 cells, 17,285 genes and 880.8 million transcripts, CellDot was the only method to complete successfully, requiring 1.8 h and 274 GB of host memory (Fig. 2d).

Together, these results demonstrate that CellDot delivers robust decontamination across diverse imaging-based ST datasets while preserving biologically meaningful expression structure and maintaining practical scalability from targeted panels to whole-transcriptome measurements. Its consistently strong performance across correction quality, marker fidelity, co-expression structure and computational efficiency supports its broad applicability to large-scale imaging-based ST analysis.

### CellDot restores genuine spatial expression patterns and enables accurate whole-transcriptome imputation

To evaluate CellDot from a spatial perspective and determine whether correction preserves biologically meaningful spatial expression patterns, we focused on the colorectal cancer dataset (Fig. 3a,b). In addition to Xenium, matched samples were profiled using the whole-transcriptome sequencing-based ST technologies Visium HD[20] and Visium[21]. Visium HD provides subcellular-resolution measurements, and its section recapitulated the major spatial organization of cell types observed in Xenium, enabling a cell-type-level comparison between the two platforms (Supplementary Fig. S10). These matched measurements served as independent evidence for validating the spatial expression patterns recovered by CellDot. Moreover, the paired scRNA-seq data provided complementary validation of cell-type expression profiles.

**Fig. 3.**
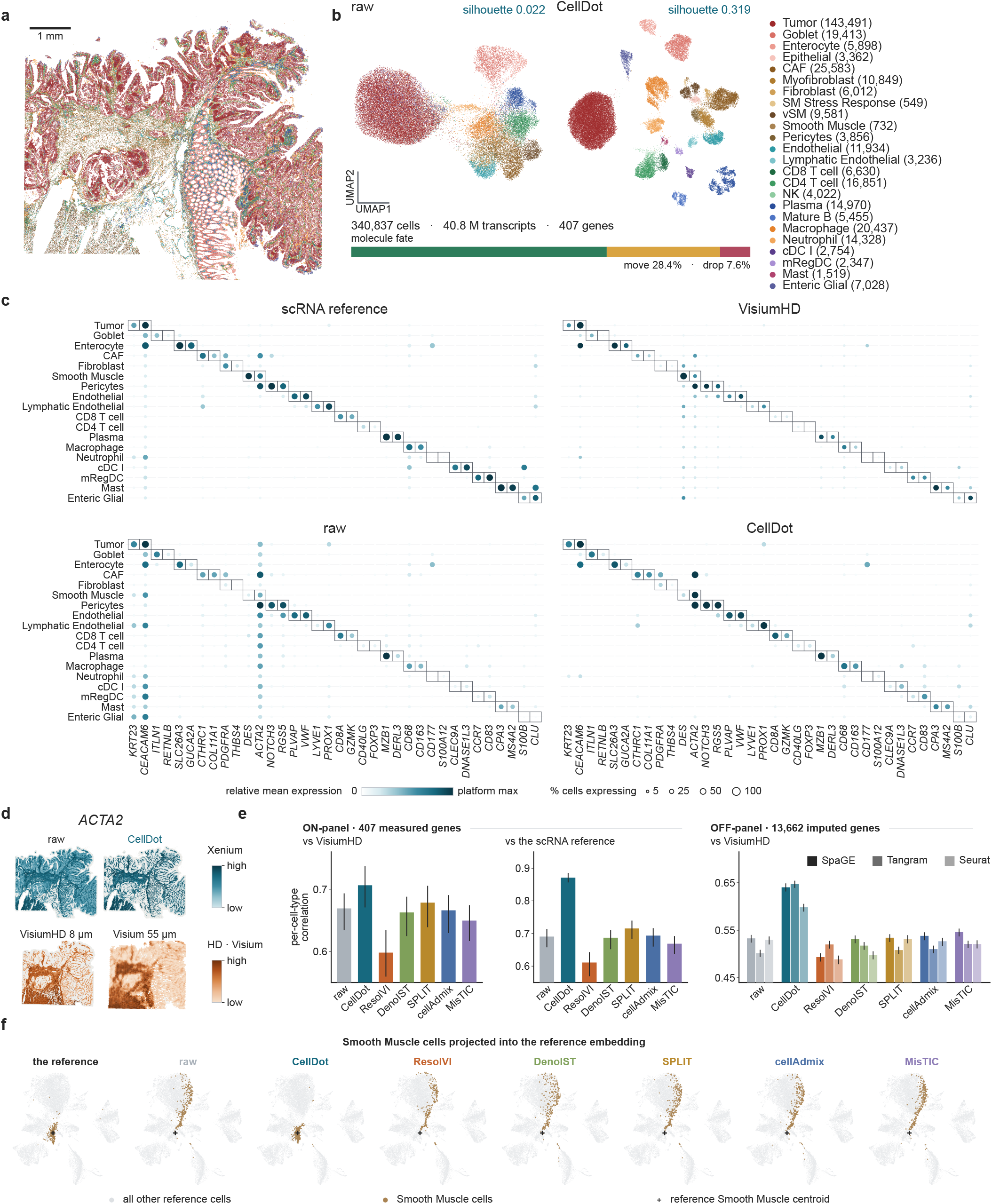
Cross-platform validation of spatial expression recovery and imputation in colorectal cancer. **a**, cell-type map of the Xenium colorectal cancer section, 340,837 cells over 24 annotated types; cell-type colors are shared with **b** and **f**. Scale bar, 1 mm. **b**, UMAP of the section before and after correction, with the average silhouette width printed above each embedding and the fate of the section’s 40.8 million transcripts summarized below. **c**, reference-derived markers, two per cell type over 18 cell types, in the single-cell reference, a serial Visium HD section, the raw Xenium counts and the CellDot results. Dot color gives mean expression relative to each platform’s maximum and dot size the percentage of cells expressing the marker; boxes mark each type’s own markers. Correction removes the off-target signal and recovers the block structure of the reference and Visium HD, raising the median on-type to off-type expression ratio from 9-fold to 34-fold. **d**, spatial expression of the smooth muscle marker *ACTA2* in the raw and corrected Xenium counts (teal) and in serial Visium HD (8 µm) and Visium (55 µm) sections of the same block (orange), each platform on its own scale. **e**, per-cell-type Pearson correlation of each layer with the independent measurements. Left and center, the 407 measured genes scored against Visium HD and against the single-cell reference; right, the 13,662 off-panel genes imputed from each layer with SpaGE, Tangram and Seurat and scored against Visium HD. Error bars, 95 % bootstrap confidence intervals over genes. **f**, smooth muscle cells of each layer projected into an embedding fitted on the reference alone and then frozen. Gray, all reference cells; cross, the reference smooth muscle centroid.

We first assessed whether CellDot restored the cell-type specificity and spatial localization of marker genes. In the raw Xenium data, several canonical markers showed widespread ectopic expression across unrelated cell types, including *KRT23, CEACAM6* and *ACTA2*. CellDot markedly reduced these off-target signals and recovered cell-type-specific expression patterns consistent with those observed in Visium HD (Fig. 3c). This improvement was also evident spatially. For instance, *ACTA2* was broadly distributed across the tumor region before correction but became restricted to smooth muscle and vascular structures after CellDot correction, closely matching its localization in both Visium HD and Visium (Fig. 3d). Similar patterns were observed across fifteen additional markers representing the major tissue compartments (Supplementary Fig. S11), supporting the recovery of biologically coherent spatial expression patterns by CellDot.

We next quantified the recovery of cell-type expression profiles using two complementary references, the scRNA-seq data and the independently measured Visium HD section. Although the scRNA-seq reference is used by CellDot, it provides a sensitive benchmark for cell-type expression, because dissociated single-cell measurements are less affected by spatial mixing and generally offer higher sensitivity and specificity than current spatial platforms[11, 28]. CellDot increased the mean correlation with the scRNA-seq reference from 0.691 to 0.871, whereas no competing method exceeded 0.715 (Fig. 3e). More importantly, CellDot also showed the highest agreement with the independent Visium HD measurements, increasing the mean correlation from 0.669 to 0.706. Improvements were observed in 11 of the 23 scorable cell types and were most pronounced for cell types with low agreement before correction (Supplementary Fig. S12).

We further asked whether recovering more faithful on-panel expression profiles could improve the inference of genes not measured by the targeted Xenium panel. Using three established imputation methods[29–31], we imputed 13,662 off-panel genes from each corrected dataset and evaluated the predictions against the whole-transcriptome Visium HD measurements. CellDot was the only method to appreciably improve imputation accuracy across the three approaches, increasing the correlation from 0.50–0.53 before correction to 0.60–0.65 after correction, whereas several competing methods reduced imputation performance (Fig. 3e and Supplementary Fig. S12).

To understand the origin of this gain, we further projected the cells of the section, under the raw counts and under each correction, into a fixed embedding fitted on the reference alone (Methods, “Projection into the reference embedding”), since imputation predicts a cell’s unmeasured genes from the reference cells it most resembles. In the raw Xenium data, contamination from the surrounding tumor caused a subset of smooth muscle cells to map into the tumor-cell region of the reference space, making them difficult to distinguish from tumor cells, with only 11% of their 50 nearest neighbors in the reference sharing the same cell-type label. After Cell-Dot correction, these cells regrouped around the smooth muscle reference population, increasing the neighbor agreement to 53% (Fig. 3f). Similar improvements were observed across most cell types, with CellDot achieving the highest neighbor agreement for 22 of the 24 cell types (Supplementary Fig. S13). These results show that accurate decontamination improves whole-transcriptome imputation by restoring cells to their appropriate transcriptional context, thereby extending the utility of targeted imaging-based ST for downstream biological analyses.

### CellDot eliminates contamination-induced cell states while preserving genuine heterogeneity

Contamination can also create spurious heterogeneity within a cell type, as transcripts from neighboring cells may generate apparent cell states or subtypes that are misinterpreted as intrinsic biological variation[12, 13, 15]. Effective correction must therefore remove these contamination-induced states without erasing genuine biological heterogeneity. To assess this, we independently sub-clustered colorectal macrophages before and after CellDot correction using Leiden clustering[32] at the same resolution. The raw data produced four sub-clusters, two of which were characterized by the non-macrophage genes *GREM1* and *PIGR*, whereas the corrected data yielded three sub-clusters (Fig. 4a).

**Fig. 4.**
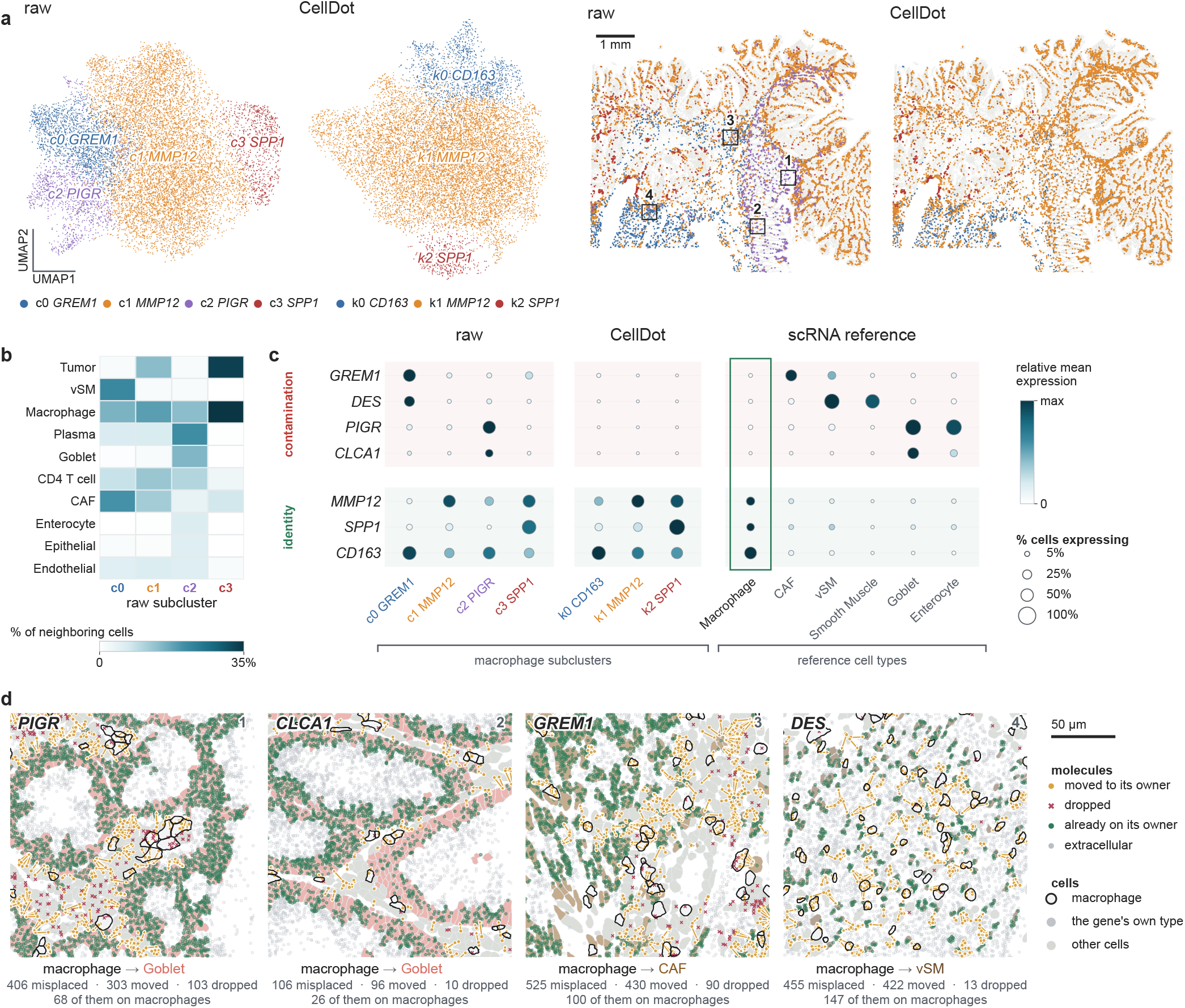
CellDot removes contamination-induced cell states while preserving genuine heterogeneity. **a**, macrophages of the colorectal cancer section, sub-clustered independently on the raw and on the corrected counts with the Leiden algorithm at resolution 0.30 and named by their top marker gene, shown in their own embeddings (left) and in their spatial distribution (right). The raw counts yielded four sub-clusters, whereas the corrected counts yielded three. The two raw sub-clusters characterized by the non-macrophage genes *GREM1* and *PIGR* had no corresponding states after correction, while the three genuine macrophage programs were preserved. Boxes 1 to 4 mark the fields shown in **d**. Scale bar, 1 mm. **b**, composition of the spatial neighborhood of each raw sub-cluster, as the percentage of neighboring cells belonging to each cell type. The *GREM1* sub-cluster sits among smooth muscle and fibroblasts, and the *PIGR* sub-cluster among goblet cells and enterocytes. **c**, the four genes that name the raw sub-clusters (contamination) and three macrophage identity genes, over the raw sub-clusters, the corrected sub-clusters and the six reference cell types that express them most. Dot color gives relative mean expression and dot size the percentage of expressing cells. The reference assigns *GREM1* and *DES* to fibroblasts and smooth muscle, *PIGR* and *CLCA1* to goblet cells and enterocytes, and none of the four to macrophages; these are the same cell types that surround the two sub-clusters in **b**. After correction the four genes are gone from the macrophage compartment and the remaining sub-clusters are named by macrophage genes. **d**, transcript fates in the four fields boxed in **a**, each corresponding to one contaminating gene. Molecules are colored by fate: retained in a cell type that expresses the gene (green), reassigned to such a cell type (gold), or discarded (red cross); macrophages are outlined in black. The numbers beneath each field report the transcripts initially detected in cell types that do not express the gene, the numbers reassigned or discarded, and the subset initially assigned to macrophages. Scale bar, 50 µm.

We traced the origin of these genes using their spatial context and the single-cell reference. The *GREM1* sub-cluster was surrounded predominantly by smooth muscle cells and fibroblasts, whereas the *PIGR* sub-cluster was enriched for neighboring goblet cells and enterocytes (Fig. 4b). Consistently, the single-cell reference showed that *GREM1* and *PIGR*, together with two other genes, *DES* and *CLCA1*, were expressed in the corresponding neighboring cell types but not in macrophages (Fig. 4c). These two sub-clusters therefore reflected contamination from the local microenvironment rather than intrinsic macrophage states.

After correction, all four contamination-associated genes were removed from macrophages, and the two artifactual sub-clusters disappeared, with their cells redistributed among three macrophage-defined sub-clusters. Importantly, this removal of spurious states did not erase genuine heterogeneity. The corrected macrophages retained three distinct programs, characterized by *C1Q*/*CD163, MMP12*/interferon and *SPP1*-*APOE* expression, respectively (Supplementary Fig. S14). These correspond to the *C1QC*^+^ and *SPP1*^+^ tumor-associated macrophage subsets described in colorectal cancer. *C1QC*^+^ macrophages are engaged in phagocytosis and antigen presentation, whereas *SPP1*^+^ macrophages promote angiogenesis and tumor progression, and the two respond differently to myeloid-targeted therapies[33]. CellDot also improved the molecular definition of these genuine states. In the raw data, 11 of the top 25 markers of the *SPP1* sub-cluster were tumor-associated genes, whereas none remained among the top markers after correction.

The molecule-level assignments produced by CellDot allow the disappearance of these spurious states to be traced directly to the underlying transcripts. For each of the four contaminating genes, transcripts detected in cell types lacking expression of that gene were predominantly reassigned or discarded, with reassigned transcripts preferentially transferred to neighboring cell types that expressed the corresponding genes (Fig. 4d). Together, these results show that CellDot eliminates contamination-induced cell states while avoiding overcorrection and preserving genuine biological heterogeneity. The resulting substructure therefore more faithfully reflects intrinsic cellular biology rather than signals acquired from neighboring cells.

### CellDot removes spurious cell-cell communication and recovers masked interactions

Imaging-based ST provides the position of every molecule, enabling a geometric validation of the transcript fate assignments. Transcripts originating from neighboring cells are expected to localize preferentially near the cell boundary and toward the putative source, whereas endogenous transcripts should be more enriched in the cell interior and show greater overlap with the nucleus[7, 9, 12]. We therefore first evaluated whether the CellDot assignments followed these geometric expectations and then examined how transcript correction affected downstream cell-cell communication inference.

On the Xenium 5K lung adenocarcinoma section (278,328 cells and 22 annotated cell types; Fig. 5a,b), the CellDot assignments followed the expected geometry in terms of nuclear overlap, distance to the nucleus and movement direction (Fig. 5c,d). Among retained transcripts, 49% overlapped a nucleus, compared with 33% of reassigned and 28% of discarded transcripts. Their mean distance to the host-cell nucleus likewise increased from 0.92 µm for retained transcripts to 1.04 µm for reassigned and 1.99 µm for discarded transcripts. In addition, 91% of reassigned transcripts were already located on the side of the host cell facing the destination cell. The same ordering was observed across all four sections (Fig. 5d), supporting the expected distinction between endogenous transcripts in the cell interior and contaminating transcripts at peripheral, source-facing locations.

**Fig. 5.**
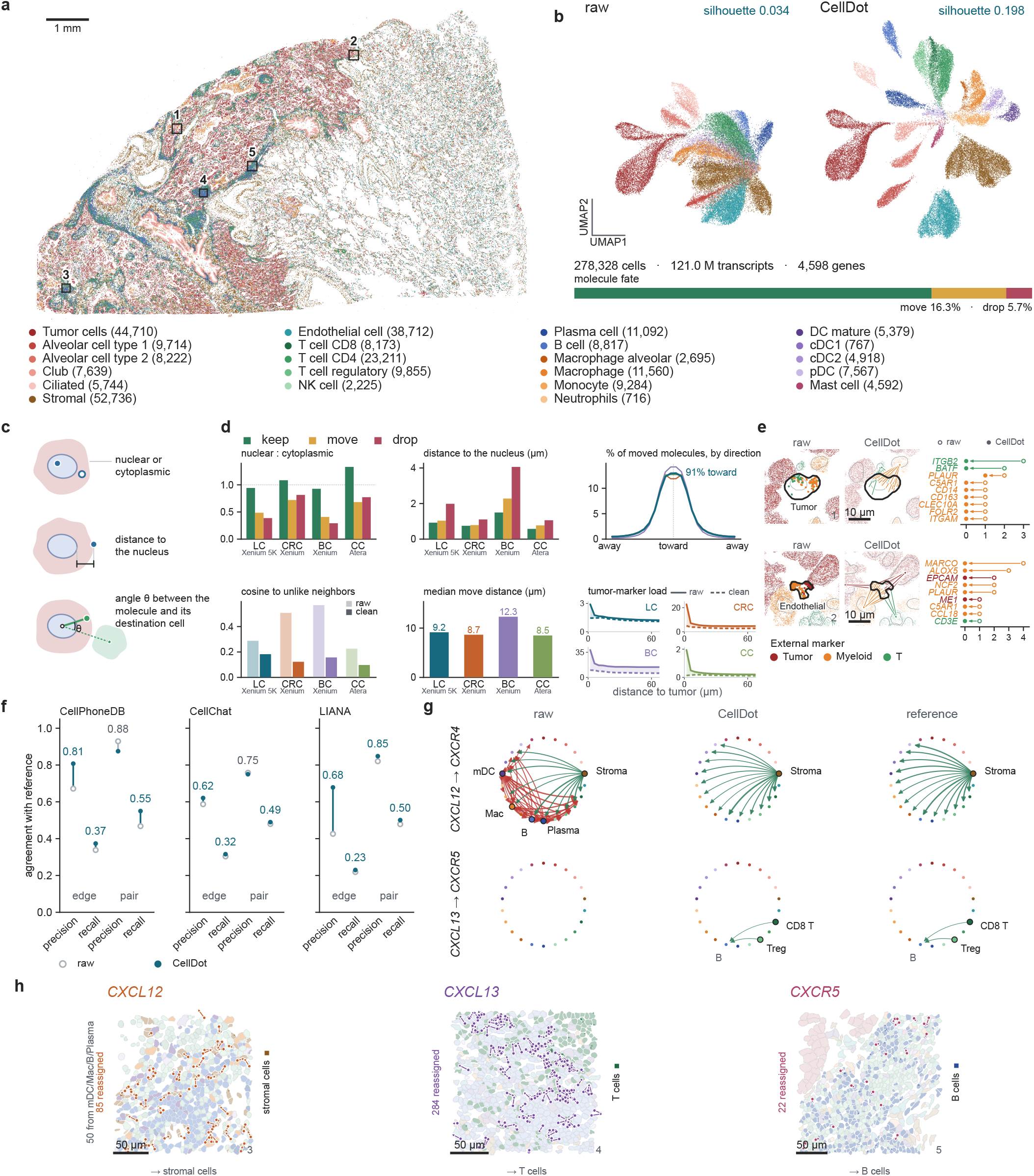
Subcellular geometry validation of transcript reassignment and improved cell-cell communication inference. **a**, cell-type map of the Xenium 5K lung adenocarcinoma section, 278,328 cells over 22 annotated types; boxes 1 to 5 mark the fields shown in **e** and **h**. Scale bar, 1 mm. **b**, UMAP before and after correction, with the average silhouette width printed above each embedding and the fate of the section’s 121.0 million transcripts summarized below. **c**, the three subcellular quantities measured in **d**, including whether a molecule lies in the nucleus or the cytoplasm, its distance to the nucleus, and the angle between the molecule and its destination cell. **d**, distribution of these features by transcript fate across all four sections (LC, lung cancer; CRC, colorectal cancer; BC, breast cancer; CC, cervical cancer). Although none of these features is used by CellDot, retained transcripts show the highest nuclear overlap and the shortest distance to the nucleus, whereas discarded transcripts show the lowest nuclear overlap and the greatest distance. Among reassigned transcripts, 91 % move toward the side of the host cell facing the destination cell, with median reassignment distances ranging from 8.5 to 12.3 µm across sections. Bottom right, mean tumor-marker abundance in non-tumor cells as a function of distance to the nearest tumor cell before (solid) and after (dashed) correction. CellDot reduces the tumor-associated expression halo and decreases the cosine similarity between neighboring cells of different types across all sections. **e**, two representative single cells corresponding to boxes 1 and 2 in **a**, shown before and after correction. Cells in each field belong to the tumor, myeloid or T-cell compartment. Molecules are colored by the compartment of the cell to which they are assigned, whereas molecules reassigned from the focal cell are colored by the compartment of their destination. Right, expression of markers from the other two compartments in each focal cell, with open circles indicating raw counts, filled circles corrected counts and arrows the reduction after correction. *ITGB2, EPCAM* and *ME1* are included although their specificity in the reference falls slightly below the threshold used to select the other markers. Discarded molecules (49 and 18) and within-compartment reassignments (59 and 0), which arise where the boundaries of same-type cells overlap, are not shown. Scale bars, 10 µm. **f**, agreement of ligand-receptor inference from the raw and the corrected counts with the same inference on the single-cell reference, for CellPhoneDB, CellChat and LIANA, at two levels: the ligand-receptor pair, and the edge, which adds the sending and receiving cell types. Open circles, raw; filled circles, CellDot. Precision is agreement with the reference’s calls and recall is coverage of them; every edge-level measure rises after correction. **g**, two ligand-receptor pairs as sender-to-receiver networks over the 22 cell types, for the raw counts, CellDot and the reference. The raw counts report five sender types for *CXCL12*, of which the reference supports only stroma; correction removes the four false senders (mature dendritic cells, B cells, plasma cells and macrophages) and recovers the two *CXCL13*-*CXCR5* senders the raw counts miss. **h**, the molecules behind **g** in three fields boxed 3 to 5 in **a**, with an arrow from each reassigned transcript’s detected position to its assigned cell. In the *CXCL12* field the four false sender types are filled in their own colors; they hold 260 of the field’s 552 cells and supply 50 of its 85 reassigned transcripts. The reassigned *CXCL12, CXCL13* and *CXCR5* transcripts move onto stromal cells, T cells and B cells, respectively. Scale bars, 50 µm.

Correction also reduced the contamination-driven similarity between adjacent cells. In the lung section, the cosine similarity between neighboring cells of different types decreased from 0.29 to 0.18, and the elevated tumor-marker signal in non-tumor cells near the tumor boundary was markedly reduced (Fig. 5d). This was evident at the single-cell level (Fig. 5e). In a tumor cell surrounded by immune cells, *CD14* and *CD163* localized near the boundary facing neighboring myeloid cells, while *ITGB2* localized toward an adjacent T cell; CellDot reassigned these transcripts to the corresponding neighboring cells and removed all three signals from the tumor cell. Similarly, in an endothelial cell adjacent to the tumor mass, the tumor markers *EPCAM* and *ME1* were completely removed after correction.

These misplaced transcripts can directly distort cell-cell communication inference by assigning ligands or receptors to the wrong cell types. We therefore applied CellPhoneDB, CellChat and LIANA[34–36] to the raw and corrected counts and compared the inferred interactions with those obtained from the matched single-cell reference, requiring agreement in the ligand-receptor pair as well as in the sender and receiver cell types (Fig. 5f). CellDot generally improved the agreement across all three methods.

For example, in the raw data, *CXCL12* was inferred to originate from five cell types, four of which were not recognized as senders in the reference, including mature dendritic cells, B cells, plasma cells and macrophages. After correction, only the stromal source remained, consistent with both the reference and the established role of *CXCL12* as a stromal fibroblast-derived chemokine acting on *CXCR4*^+^ immune cells[37]. Conversely, the *CXCL13*-*CXCR5* interaction was missed in the raw data but recovered after correction with the appropriate Tcell senders (Fig. 5g), consistent with *CXCL13*-producing T cells recruiting *CXCR5*^+^ B cells in human lung cancer[38].

At the molecule level, CellDot reassigned *CXCL12* transcripts from immune cells to stromal and endothelial cells, thereby removing the false senders (Fig. 5h). Similarly, *CXCL13* transcripts were reassigned from B cells and dendritic cells to T cells, while *CXCR5* transcripts moved from T cells to B cells, restoring the missing interaction by correcting both the sender and the receiver assignments. Across the whole section, 98% of the transcripts of these four genes were retained, while the number of cells carrying them decreased by 23%, concentrating their expression within the appropriate cell types (Supplementary Fig. S15). A similar restoration of sender-receiver assignments was observed for additional ligand-receptor pairs (Supplementary Fig. S16). Together, these results show that CellDot corrects transcript-level misassignment in a manner supported by sub-cellular geometry, removing spurious communication signals while recovering interactions that were previously obscured by contamination.

### CellDot preserves malignant cell states and improves spatial niche analysis at whole-transcriptome scale

We finally evaluated CellDot on the whole-transcriptome cervical cancer section, for which no baseline method was computationally applicable. CellDot processed 880.8 million transcripts in 1.8 hours, reassigning 24% and discarding 8% (Fig. 6a,b). We examined whether correction at this scale preserves genuine cell states and remains effective for genes beyond the marker-focused panels of earlier targeted platforms.

**Fig. 6.**
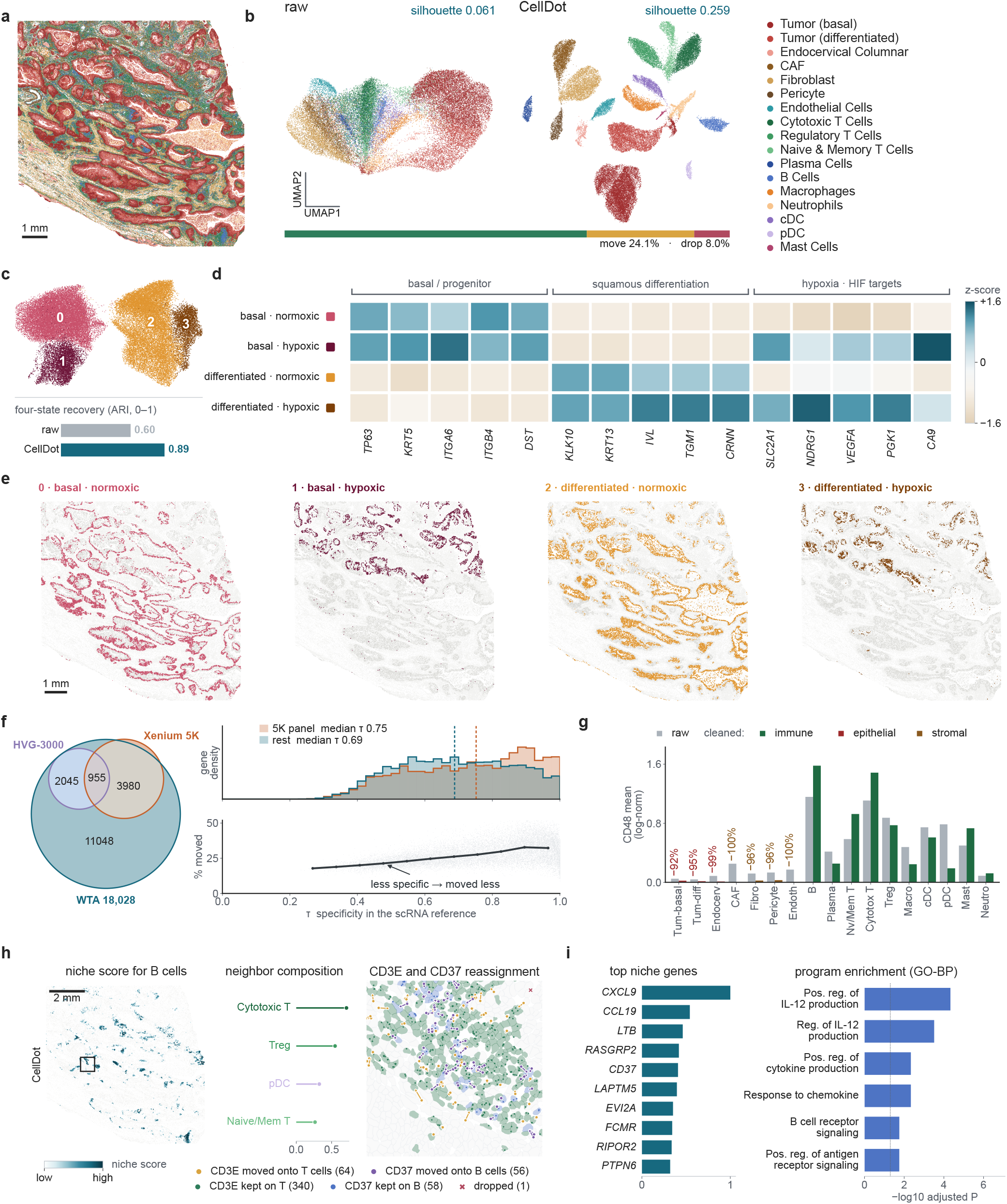
Whole-transcriptome decontamination preserves malignant cell states and restores spatial niche programs in cervical cancer. **a**, cell-type map of the whole-transcriptome cervical cancer section, 717,576 cells over 17 annotated types. Scale bar, 1 mm. **b**, UMAP before and after correction, with the silhouette width printed above each embedding and the fate of the section’s 880.8 million transcripts over 17,285 genes summarized below. **c**, malignant cells of the corrected layer sub-clustered into four states, and the adjusted Rand index (ARI) of each layer’s sub-clustering against a fixed label that crosses this study’s lineage call with the data release’s hypoxic group (0.60 raw, 0.89 CellDot). **d**, the four states over the 15 marker genes that define them, in three modules: basal/progenitor, squamous differentiation and hypoxia. The states organize along two axes, basal to differentiated and normoxic to hypoxic. **e**, the four states in space, one highlighted per tile; the two hypoxic states occupy the same upper region of the section. Scale bar, 1 mm. **f**, left, overlap of the 18,028 genes profiled by Atera with the 5,101-gene Xenium 5K panel (the 5,001-gene Xenium Prime 5K panel plus a 100-gene add-on) and the 3,000 highly variable genes of the reference; 11,048 genes are in neither. Top right, the distribution of cell-type specificity *τ* in the reference for panel genes and for the rest (median 0.75 against 0.69), showing that the genes no panel carries are expressed more broadly. Bottom, the percentage of each gene’s transcripts reassigned against *τ*: the less specific a gene, the less of it is moved. **g**, mean *CD48* expression per cell type in the raw counts (gray) and after correction (colored by compartment), with the percentage removed printed over each non-immune type. *CD48* is absent from the 5K panel and marks the immune lineage as a whole; correction removes 92 to 100 % of the non-immune signal while the immune types keep theirs. Where removal rounds to 100 % no corrected bar is drawn; the remaining corrected values lie far below the axis and are drawn at an expanded scale that preserves their order. **h**, the T-cell-adjacent B-cell niche fitted with NicheScope on the corrected counts. Left, the niche score in space, with the field of the molecule map marked; the mark is drawn larger than the 180 µm field so that it remains visible. Scale bar, 2 mm. Middle, the fitted neighbor weights of the four cell types the niche is built on, a T-cell and dendritic-cell neighborhood around B cells whose composition is consistent with a tertiary lymphoid structure. Right, *CD3E* and *CD37* transcripts in that field, drawn by fate: 64 *CD3E* transcripts move onto T cells (gold), 16 of them off B cells, and 340 stay on T cells (green); 56 *CD37* transcripts move onto B cells (violet), 44 of them off T cells, and 58 stay on B cells (blue); one *CD3E* transcript is dropped. T and B cells are filled in their own colors, and transcripts on other cell types or assigned to no cell are not drawn. The two genes cross the same boundary in opposite directions. **i**, the ten leading genes of the corrected niche program, ranked by the loading they carry in it, and six of the nine pathways the program enriches for (GO biological process, −log10 adjusted *P*). Seven of the ten genes carry no loading in the raw fit and are recovered by correction. *CXCL9* and *CCL19* load as strongly on the raw fit as on the corrected one and are assigned by the reference to macrophages and stroma; they describe the neighborhood the niche sits in rather than an effect of correction. Fitted on the raw counts, the same niche keeps the same neighborhood but is built from T-cell genes and enriches for T-cell receptor signaling (Supplementary Fig. S21).

To determine whether CellDot preserves genuine intratumoral heterogeneity, we sub-clustered the malignant compartment and identified four states organized along two axes, basal to differentiated and normoxic to hypoxic (Fig. 6c,d). The two hypoxic states occupied the same tumor region, supporting the spatial coherence of the hypoxic axis (Fig. 6e). We further compared the raw and corrected sub-clusters with a fixed four-class annotation derived from the expert hypoxic labels released with the dataset[39] and the malignant lineage identity. The adjusted Rand index increased from 0.60 to 0.89 after correction, indicating that CellDot sharpened rather than erased these states. This agreement extended beyond malignant cells. The corrected profiles formed coherent groups according to the 26 expert-defined annotations (Supplementary Fig. S17). Consistently, CellDot reduced the ectopic expression of expert-selected and reference-derived markers while preserving their expression within the corresponding cell groups (Supplementary Figs. S18 and S19).

We next examined how CellDot performs on genes not included in targeted imaging-based ST panels. Such panels typically measure a predefined set of genes enriched for cell-type markers, leaving many genes with broader or less cell-type-specific expression unmeasured. Of the 18,028 genes profiled by Atera, 11,048 were absent from both the 5,101-gene Xenium 5K panel (the 5,001-gene Xenium Prime 5K panel with the 100-gene add-on of a cervical cancer Xenium dataset) and the 3,000 most variable genes of the reference. These genes were, on average, less cell-type specific and ranged from lineage-level markers to broadly expressed housekeeping genes (Fig. 6f), where the cell-type specificity score *τ* is calculated based on Yanai et al.[40], as detailed in the Methods. Because CellDot assigns a fate to every transcript within a single optimal-transport problem, it requires no prior gene selection and corrects all genes simultaneously. Genes represented in the targeted sets and those outside them showed similar overall correction rates, while within both groups the reassignment rate increased with cell-type specificity (Fig. 6f). As an example, *CD48* is absent from the Xenium 5K panel and marks the immune lineage broadly rather than any individual immune cell type. In the raw data, *CD48* was detected across all non-immune cell types. CellDot removed 92–100% of this off-lineage signal, restoring near-zero expression in non-immune populations while preserving expression in immune cells (Fig. 6g). At the molecule level, *CD48* transcripts assigned to non-immune cells were predominantly reassigned to immune cells or discarded, whereas most transcripts already assigned to immune cells were retained (Supplementary Fig. S20). These results show that whole-transcriptome correction extends beyond conventional cell-type markers to genes with broader, lineage-level expression patterns.

Finally, we examined spatial niche programs. Spatial niches are multicellular neighborhoods in which the expression state of a cell is associated with the composition of cell types around it, and a niche program is the set of genes that vary with this neighborhood composition. Such programs are particularly susceptible to contamination, because both genuine niche effects and contamination can appear as expression changes associated with neighboring cells. NicheScope[41] identifies niches and their programs by coupling each cell’s own expression with the cell-type composition of its neighborhood, returning for each niche a neighbor cell-type profile together with the associated gene program. We applied NicheScope independently to the raw and corrected counts. Both analyses identified the same B-cell niche in the same spatial locations, surrounded by cytotoxic and regulatory T cells and dendritic cells (Fig. 6h and Supplementary Fig. S21). This cellular composition, together with *CCL19, CXCL9* and the recovered lymphotoxin gene *LTB*, was consistent with a tertiary lymphoid structure associated with the response to immunotherapy[42].

Despite identifying the same niche, the raw and corrected data yielded markedly different niche-associated expression programs. In the raw counts, the program was dominated by T-cell genes including *CD3E, TRAC* and *GZMK* and enriched for T-cell receptor signaling, incorrectly suggesting a T-cell-like program in B cells (Supplementary Fig. S21). After correction, the program instead contained B-cell and leukocyte genes such as *CD37* and *LTB* and was enriched for B-cell receptor signaling and cytokine production (Fig. 6i). At the molecule level, all 16 *CD3E* transcripts assigned to B cells in one field were reassigned to T cells, while 44 *CD37* transcripts were reassigned from T cells to B cells (Fig. 6h). A second B-cell niche adjacent to tumor cells showed the same pattern (Supplementary Fig. S22). Together, these results show that CellDot enables reliable whole-transcriptome correction while preserving genuine tumor states, correcting genes not included in targeted panels, and improving the biological fidelity of spatial niche analysis.

## Discussion

In this work, we presented CellDot, a decontamination method for imaging-based ST that treats contamination as an assignment problem over individual molecules. Each detected transcript is kept in its segmented host cell, reassigned to a nearby cell or discarded as background, and all fates are resolved jointly in a single capacitated optimal-transport problem guided by a paired single-cell reference. Across three Xenium tumor sections, Cell-Dot recovered cell-type structure better than all baseline methods, without relying on extensive transcript deletion to achieve superior performance. On a whole-transcriptome Atera section, CellDot was the only applicable method. Matched Visium HD and Visium measurements further confirmed that the corrected counts recovered cell-type expression profiles and spatial marker patterns, while the inferred transcript fates were independently supported by subcellular geometry. In downstream analyses, CellDot eliminated contamination-induced macrophage states while preserving genuine heterogeneity, recovered the correct senders of cell-cell communication, and restored biologically coherent programs of a tertiary lymphoid niche at whole-transcriptome scale.

Three key design principles enable the superior performance of CellDot. First, CellDot operates at the level of individual molecules, allowing contaminating transcripts to be reassigned rather than simply removed. A transcript can be transferred to a specific neighboring cell, so correction can both eliminate false signals and recover missing ones, as illustrated by the restoration of previously obscured cell-cell communication patterns. Second, CellDot estimates the key quantities directly from the data, including gene-specific background levels, platform scaling factors and capacity margins. This data-adaptive design improves correction accuracy across datasets while eliminating the need to manually specify a contamination rate. Third, the transport problem is solved tile by tile with a computational cost that grows linearly with the number of transcripts. Consequently, the same framework can be applied across datasets ranging from a 313-gene targeted panel to whole-transcriptome measurements.

However, CellDot has limitations that warrant further improvement. First, CellDot assumes that although imaging-based ST data contain contamination, only a minority of transcripts within most cells are affected, such that cell identity remains largely preserved and reliable cell-type annotations can still be obtained. These annotations, together with a matched single-cell reference, provide the basis for the subsequent decontamination. While this assumption is expected to hold for most cells, low-count cells or regions with unusually severe contamination may still be misannotated and consequently corrected toward an inappropriate reference profile. Reference quality is equally important. A reference that is itself noisy or lacks cell types present in the tissue cannot provide an appropriate prior for those cells and may limit correction accuracy. A patient-matched reference is preferable when available, but is not essential, as demonstrated by the lung cancer analysis, which used a public atlas reference and still performed well. Second, CellDot estimates gene-specific background levels from transcripts detected in extracellular regions. Tissues with very limited extracellular space therefore provide less information for this estimate and may lead to less reliable background quantification. The estimate also depends on segmentation quality. Although nuclei can generally be delineated with high precision, cell boundaries may remain more uncertain[43]. CellDot mitigates moderate boundary errors by dilating cell boundaries before defining extracellular transcripts, but substantial segmentation inaccuracies can still bias the estimated background.

Several extensions may further broaden the utility of CellDot. Since imaging-based ST platforms already record the *z* position of each transcript, the same transport framework can be extended naturally from two to three dimensions without a change of principle. The modular cost function can likewise incorporate additional evidence, such as segmentation confidence, so that correction can account for uncertainty in cell boundaries rather than treating segmentation as fixed. In addition, the current cost evaluates each gene independently. Incorporating gene-gene co-expression structure from the reference could encourage assignments that preserve coordinated expression programs, thereby improving the recovery of coherent within-cell expression patterns.

As imaging-based ST expands toward whole-transcriptome profiling at increasingly large scale, the accurate assignment of individual transcripts to their cellular origins becomes increasingly important for faithful biological interpretation. CellDot provides a principled, traceable and scalable framework for molecule-level decontamination, assigning each transcript an explicit fate while preserving biologically meaningful signals. Such molecule-level correction may become a prerequisite for the faithful interpretation of cellular states, tissue organization and intercellular interactions.

## Methods

### The model of CellDot

Imaging-based ST technologies detect individual transcripts together with their spatial coordinates, while cell boundaries are defined through a separate segmentation step. Specifically, in this study, the cell boundaries and the initial transcript-to-cell assignment are the standard outputs of the platform’s own analysis pipeline for each dataset (Xenium Onboard Analysis, with nucleus-expansion or multimodal stain-based cell segmentation as released with the data). These initial assignments can nevertheless be incorrect when detected transcripts derive from neighboring or spatially overlapping cells, or represent extracellular background[10–12]. CellDot addresses this problem at the molecule level by jointly determining whether each transcript should remain with its current cell, be reassigned to a nearby cell, or be attributed to the background through a capacitated entropic optimal-transport formulation[16, 17, 19].

#### Notation

We denote by *G* the set of genes shared between the spatial panel and the scRNA-seq reference, and let *G* = |*G*| be the number of shared genes. Among quality-controlled transcripts whose genes belong to *G*, those assigned by the segmentation to a host cell with a cell-type label are indexed by *m* = 1, …, *M*. Each transcript *m* has position **x**_*m*_, gene identity *g*(*m*), and host cell *h*(*m*). Cells are indexed by *k* = 1, …, *N*, where *N* is the number of segmented cells. Cell *k* has segmented area *A*_*k*_, cell type *t*_*k*_ and raw library size *N*_*k*_, defined as the number of transcripts from genes in *G* that are assigned to cell *k* by the segmentation. Transcripts of genes in *G* that are not assigned to any segmented cell constitute the extracellular pool. Those retained after the segmentation-aware preprocessing described below define the gene-specific background and are not themselves reassigned (Supplementary Note 2).

#### Reference-guided prior with platform calibration

Cell-type labels of ST datasets are obtained upstream (see more details in the “Cell-type annotation” section). For the single-cell reference, each cell is normalized to a composition over *G*, that is, its counts are divided by their sum over *G*, and these compositions are averaged within each cell type to define the reference-guided expression prior 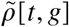, with 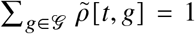 for every cell type *t*. Here, 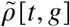 represents the probability that a transcript originating from a type-*t* cell corresponds to gene *g*. To account for systematic gene-specific differences between the single-cell reference and the spatial measurement, we estimate a platform factor *γ*_*g*_ as the ratio of the observed spatial pseudobulk abundance of gene *g* to that expected under the reference prior, following the treatment of platform effects in RCTD[26]. The factor *γ*_*g*_ rescales the overall abundance of each gene while preserving its relative expression across cell types, yielding the calibrated prior 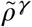, which is used only within the model, specifically in the assignment costs and the expression capacities defined below (Supplementary Fig. S1 and Supplementary Note 1).

#### Estimation of the extracellular background

CellDot uses extracellular transcripts as an empirical measurement of the contamination background. Let *n*_*g*_ denote the number of extracellular transcripts of gene *g*, and let *A*_extra_ denote the cell-free tissue area, obtained by subtracting the area occupied by segmented cells from a transcript-density-based tissue mask (Supplementary Note 2). We define

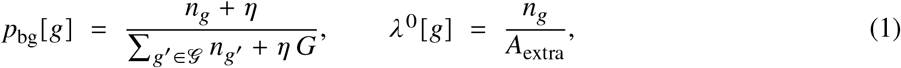

with a small pseudocount *η* = 10^−3^. These two quantities characterize complementary aspects of the background: *p*_bg_[*g*] is the probability that a background transcript corresponds to gene *g*, so that *p*_bg_ describes the gene composition of the background, whereas *λ*^0^[*g*] is the extracellular density of gene *g* in molecules per µm^2^. Both are estimated independently for each tissue section (Supplementary Fig. S2). For segmentations with tight cell boundaries, genuinely cellular transcripts immediately outside the segmented boundary can otherwise be misclassified as extracellular and inflate the background estimate. We therefore exclude extracellular transcripts within Δ = 3 µm of a cell boundary where applicable (Supplementary Note 2).

In the transport model, *p*_bg_ defines the gene composition of the background component. Like each row of 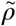, it is a unit-sum distribution over the gene panel, but it is estimated directly from the spatial section (equation 1) rather than from the single-cell reference. This places the background and the candidate cells on a comparable scale when evaluating the possible fates of each transcript, while *λ*^0^ determines how much transcript mass can plausibly be assigned to the background.

#### Candidate cells and assignment costs

Each transcript is assigned to one of a limited set of nearby candidate cells or to the extracellular background. For transcript *m*, the candidate set *N*(*m*) comprises up to *K* = 14 cells whose centroids are nearest to the transcript and lie within *R* = 15 µm of its location. This set typically includes the host cell *h*(*m*) assigned by the initial segmentation together with neighboring cells. In addition, every transcript may be assigned to a shared background destination, denoted by ∅. The cost of assigning transcript *m* to a candidate cell *k* ∈ *N*(*m*) is defined as

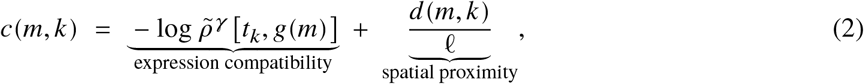

where *d*(*m, k*) is the distance from transcript *m* to the surface of cell *k*, with the cell treated as a disk of its own area (Supplementary Note 1), and *ℓ* = 4.7 µm is the distance scale. The cost of assigning a transcript to a candidate cell combines expression compatibility and spatial proximity. The expression term is the negative log-probability that a cell of type *t*_*k*_ produces a transcript of gene *g*(*m*), so it favors cell types in which the detected gene is more likely to be expressed, whereas the distance term favors cells located closer to the transcript. The cost associated with the background sink is determined solely by the gene-specific background composition. Therefore, it can be defined as follows:

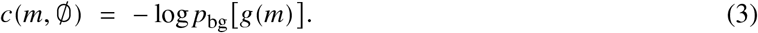

The distance term is omitted because the background reflects a spatially diffuse extracellular component rather than a localized cellular source. Consequently, genes that are abundant in the extracellular pool incur minimal costs when attributed to the background.

#### Capacity constraints

The capacity constraints limit the amount of transcript mass that can be assigned to each destination and thereby couple the individual transcript assignments into a joint optimization problem. For every cell *k* and gene *g*, we define an expression capacity and a background capacity,

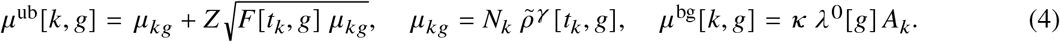

The expression capacity *μ*^ub^[*k, g*] sets an upper bound on the number of gene-*g* transcripts that cell *k* can contain after correction. Its baseline *μ*_*kg*_ is the expected number of gene-*g* transcripts in cell *k*, given the cell’s raw library size *N*_*k*_ and the calibrated reference profile of its cell type. Because expression varies among cells of the same type, constraining every cell to this expectation would remove genuine high-expression signals. We therefore add a dispersion-aware margin based on the Fano factor *F*[*t*_*k*_, *g*], defined as the variance-to-mean ratio of the raw gene-*g* counts across the spatial cells of type *t*_*k*_ . For an expected count *μ*_*kg*_, this gives a variance of *F*[*t*_*k*_, *g*] *μ*_*kg*_, and we define the upper expression capacity as the expected count plus *Z* standard deviations. We use *Z* = 2, with *F* estimated from the raw counts of each dataset and clipped to the range [1, 50]. Genes with greater within-type dispersion receive a wider allowable range, whereas genes with negligible expected expression retain a near-zero capacity, limiting the accumulation of incompatible transcripts.

The background capacity *μ*^bg^[*k, g*] limits the number of gene-*g* transcripts from cell *k* that can be assigned to the background. It is the expected number of background molecules of gene *g* within the area of cell *k*, given the measured extracellular density *λ*^0^[*g*], scaled by a factor *κ*. Rather than using a user-specified contamination rate, this capacity is determined jointly by the gene-specific background density and the cell area, allowing it to adapt to the ST datasets. Genes that are abundant in the extracellular background permit greater removal, genes that are rarely observed there permit little removal, and larger cells receive proportionally larger background capacities. We set *κ* = 1.2 for all datasets.

#### Optimal-transport formulation

Combining the assignment costs and capacity constraints defined above, Cell-Dot solves for a transport plan *P* with non-negative entries. The entry *P*_*mk*_ is the fraction of transcript *m* assigned to candidate cell *k* ∈ *N*(*m*), and *P*_*m*∅_ is the fraction assigned to the background. A transcript can therefore only be placed in one of its own candidate cells or in the background. Writing *C* for the matrix of assignment costs, with entries *C*_*mk*_ = *c*(*m, k*) and *C*_*m*∅_ = *c*(*m*, ∅), the plan is the solution of

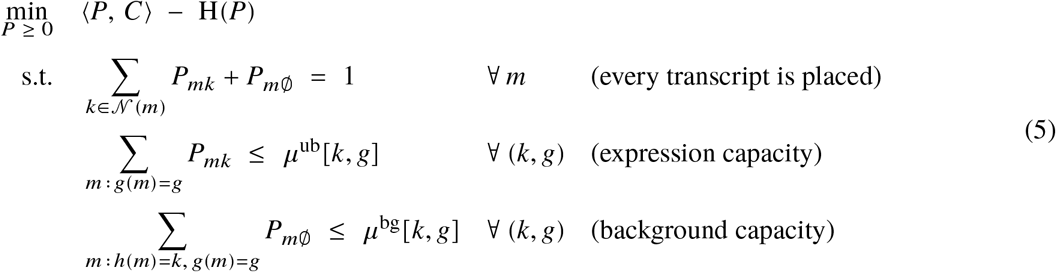

where ⟨*P, C*⟩ = ∑_*m*_ ∑_*k*_ *P*_*mk*_ *C*_*mk*_ is the total assignment cost, H(*P*) = − ∑_*m*_ ∑_*k*_ *P*_*mk*_ (log *P*_*mk*_ −1) is the entropy of the plan, and both sums run over the destinations *k* ∈ *N*(*m*) ∪ {∅} of each transcript. The first constraint, ∑_*k*∈*N*(*m*)_ *P*_*mk*_ +*P*_*m*∅_ = 1, states that the unit mass of each transcript *m* is distributed over its candidate cells and the background, so that no transcript is lost or duplicated. The second constraint, ∑_*m*∶*g*(*m*)=*g*_ *P*_*mk*_ ≤ *μ*^ub^[*k, g*], sums the mass of all gene-*g* transcripts that list cell *k* among their candidates, whether they are currently hosted by cell *k* or by one of its neighbors, and bounds this mass by the expression capacity of cell *k*. It thereby limits the number of gene-*g* transcripts that cell *k* can retain or receive. The third constraint, ∑_*m*∶*h*(*m*)=*k, g*(*m*)=*g*_ *P*_*m*∅_ ≤ *μ*^bg^[*k, g*], sums the mass that the gene-*g* transcripts hosted by cell *k* send to the background and bounds it by the background capacity of cell *k*, so that the number of gene-*g* transcripts removed from cell *k* cannot exceed *μ*^bg^[*k, g*]. Both capacities are upper bounds rather than target values, restricting implausible assignments without forcing reassignment or removal. Every unit of transported mass is an observed transcript that is retained, reassigned or assigned to the background, so CellDot does not introduce unobserved molecules, and every change in the corrected expression profile can be traced back to individual transcripts and their spatial locations. The entropy term makes the objective strictly convex, so that the transport plan is unique and can be computed efficiently at the scale of whole tissue sections by the iterative scaling procedure described below.

#### Transport optimization and transcript fate assignment

Equation (5) is solved using Sinkhorn-type alternating projections[17, 18]. The background capacity is enforced as a hard constraint, whereas the expression capacity is treated as a soft constraint, because it represents a type-level expression expectation applied to individual cells (Supplementary Note 3). The resulting transport plan is soft, that is, each transcript’s unit mass can be split across its candidate destinations. CellDot converts this plan into discrete transcript fates according to the transport preferences while reconciling the background assignments with locally pooled background capacities. Because the cell-level background capacities are often smaller than one molecule, the background budget is aggregated within a sliding 100 µm window so that the discrete removals remain consistent with the locally estimated background (Supplementary Note 4).

#### Implementation and scalability

Tissue sections are processed in 500 µm tiles with a 15 µm halo matching the assignment radius. Molecules within the halo participate in the transport optimization, whereas assignments are recorded only for molecules in the tile core, ensuring that each molecule receives a single final fate. Run time scales linearly with the number of molecules, while memory is bounded by the tile size rather than by the section size. CellDot is implemented in Python using PyTorch[44], NumPy[45] and SciPy[46]. On the largest section analyzed here, comprising 717,576 cells and 17,285 genes, with 880.8 million of the 1.24 billion detected transcripts entering the solver, CellDot finished in 1.8 h on a single GPU with a peak host memory of 274 GB. Each run outputs the corrected expression profiles together with a molecule-level fate table, allowing every correction to be traced to the underlying transcript assignment. The same parameter settings were used across all datasets in this study (Supplementary Table 1).

### Cell-type annotation

Each dataset was annotated before correction, and the same cell-type labels were provided to all methods requiring them and used consistently for downstream evaluation. Cell-type labels were transferred from the corresponding single-cell reference using scANVI[47, 48], which was trained on the reference and the spatial data together and returned a predicted label and posterior probability for each spatial cell (Supplementary Note 5). All genes shared between the spatial data and the reference were used as input to the model for every dataset.

### Benchmarking analysis

CellDot was compared with five baseline methods, including resolVI[13], MisTIC[15], DenoIST[14], SPLIT[11] and cellAdmix[12]. All methods were run using their published default settings (Supplementary Note 6), with the same cell-type annotations provided whenever required. Cells that could not be processed by a given method, for instance because they did not meet its minimum-count requirement, retained their raw counts. This ensured that all datasets remained complete and that apparent transcript removal was not introduced by excluding unprocessed cells.

#### resolVI

resolVI learns a latent representation of each cell and generates corrected counts from it. We used resolVI as implemented in scvi-tools version 1.4.3, run in semi-supervised mode so that it receives the same cell-type labels as the other methods, trained for the default 50 epochs. Corrected counts were taken as the median of 25 posterior samples; cells with fewer than five counts, which the model cannot process, were kept as raw.

#### MisTIC

MisTIC reassigns and removes individual transcripts using a neural classifier over each molecule’s local context. We used MisTIC version 0.0.2 with its default neighborhood size and its default reassignment and removal threshold grids, trained under the package’s built-in convergence test. Its removal mode was set to best because the default conservative setting disables removal entirely.

#### DenoIST

DenoIST uses a Poisson mixture model with local-neighborhood information to infer, for each cell– gene pair, whether the observed gene counts are likely to represent endogenous expression or contamination. Model parameters are estimated by expectation maximization, and gene counts classified as contamination according to the posterior probability are removed. We used DenoIST version 1.1.0 with all parameters at their defaults (neighborhood distance 50 µm, 200 bins, posterior cutoff 0.6, 10 initializations). DenoIST operates without cell-type labels.

#### SPLIT

SPLIT purifies each cell’s expression profile based on the deconvolution results obtained through RCTD with a single-cell reference, for which we provided the same reference used by CellDot. We ran SPLIT v0.3.0 following the published pipeline and RCTD settings, with the residual-contamination flag set to its default value.

#### cellAdmix

cellAdmix (v0.1.0) decomposes local expression profiles into non-negative matrix factorization components and classifies each component as native or foreign, with the molecules assigned to foreign components removed during correction. The number of components was determined according to the rule described in the cellAdmix paper, as 1.5 times the number of cell types with rounding, while all other parameters followed the published workflow. cellAdmix does not require a single-cell reference.

### Evaluation metrics

#### Cell-type separability

Separability is measured by the average silhouette width[49] on the shared cell-type labels. The evaluation space was defined once from the raw data and reused for all corrected datasets (2,000 highly variable genes, cells with at least 20 raw counts, 30 principal components).

#### Residual contamination

Residual contamination was quantified using three marker-based metrics. The off-type marker fraction measures the proportion of marker-gene transcripts detected in cell types that do not express the corresponding marker. We additionally computed the mutually exclusive co-expression rate (MECR) and the positive marker purity (PMP)[14]. All marker sets were derived exclusively from the single-cell reference and not from the spatial data being evaluated. For the off-type marker fraction, we used the two most specific reference markers for each cell type, whereas the markers for MECR and PMP were selected following the differential-expression procedure described in DenoIST.

#### Over-correction

Because improved separability can also result from excessive transcript removal, we report the deletion fraction alongside the performance metrics as a diagnostic rather than a score. We additionally report the number of genes detected per cell, which decreases under overcorrection[11].

#### Cross-platform concordance

On the colorectal section, each layer is additionally scored by the Pearson correlation of its per-cell-type expression profiles against the matched single-cell reference and against Visium HD, as described under “Cross-platform validation on the colorectal section”.

### Computational efficiency

Runtime and peak memory were evaluated using nested subsets of cells and genes sampled from the whole-transcriptome cervical cancer section, such that each larger benchmark contained all cells and genes of the preceding smaller one. All methods were run on the same server, equipped with two 22-core CPUs, 754 GB of memory and NVIDIA Tesla V100 (16 GB) GPUs. Run time was fitted as *T* ∝ *N*^*β*^*G*^*α*^, where *N* and *G* denote the numbers of cells and genes, respectively. The scaling with cell numbers and with gene numbers was fitted separately, and extrapolation to the full section was based on the slopes estimated at the largest benchmark sizes. Runs with a projected runtime exceeding 12 h were not executed, and the corresponding projected runtime was reported instead. Full benchmarking details are provided in Supplementary Note 7.

### Cross-platform validation on the colorectal section

The colorectal Xenium section is matched by Flex single-cell RNA-seq, Visium and Visium HD data from the same specimen. Visium HD was analyzed at the 8 µm bin level with the cell-type labels released with the dataset. To compare expression across platforms, each cell or bin was normalized to counts per ten thousand, profiles were averaged within each cell type, and each gene’s profile was then normalized to sum to one across cell types, so that the comparison asks which cell types own each gene. For every cell type present with at least 200 Xenium cells, 50 Visium HD bins and 40 reference cells (23 types), we computed the Pearson correlation between platforms across the shared panel genes and averaged the correlations over cell types; confidence intervals are bootstrapped over genes. The same procedure scored every method’s output layer.

### Gene imputation

To test whether corrected counts better support the prediction of genes outside the panel, we imputed off-panel expression from each layer with SpaGE[29] (its published GitHub implementation), Tangram[30] (v1.0.4, cluster mode, 200 epochs) and the anchor-based transfer of Seurat[31] (v5.5.1, 50 principal components). Each tool was trained on the panel genes shared with the single-cell reference and applied to 13,662 off-panel genes detected in both the reference and Visium HD, and the imputed profiles were scored against Visium HD by the same cell-type-profile correlation as above.

### Projection into the reference embedding

To ask whether correction moves each cell toward the reference cells of its own type, which are the cells that imputation borrows from, we embedded the spatial cells in a space fitted on the single-cell reference alone and then frozen (Fig. 3f and Supplementary Fig. S13). The reference was normalized to counts per ten thousand and log-transformed, restricted to the 407 panel genes shared with the Xenium section, standardized gene by gene with its own mean and standard deviation, reduced to 50 principal components and embedded with UMAP (30 neighbors, minimum distance 0.3, seed 0). The raw counts and each corrected layer were normalized and logtransformed in the same way, standardized with their own gene-wise mean and standard deviation, projected with the reference’s principal-component loadings and transformed into the fixed UMAP model, without refitting any step.

### Co-expression analysis

Gene-gene co-expression was estimated using CS-CORE[27] with default settings and the same set of cells for every layer. Marker blocks were defined from the single-cell reference: each shared panel gene was assigned to the cell type with the highest mean expression and retained if its specificity was at least 1.5-fold above the panel mean and its detection rate was at least 2%, with the top 25 genes retained for each cell type. Co-expression between genes within the same marker block reflects the preservation of cell-type-associated expression programs, whereas co-expression between different blocks reflects contamination-driven mixing. Agreement between each layer and the reference was quantified by the correlation of their co-expression estimates across all gene pairs (Supplementary Fig. S9).

### Sub-clustering of the macrophage compartment

Macrophages of the colorectal section were sub-clustered independently on the raw and the corrected counts with an identical pipeline. Counts were normalized to counts per ten thousand, log-transformed and scaled, followed by PCA with 30 components, construction of a 15-nearest-neighbor graph, and Leiden clustering[32] at a fixed resolution of 0.30. Each sub-cluster was labeled by its top-ranked differentially expressed gene, identified using a Wilcoxon test against all other sub-clusters. For each sub-cluster, the spatial neighborhood was summarized by averaging the cell-type composition of the 15 nearest tissue neighbors of its constituent cells.

### Subcellular geometry analysis

The subcellular position of each transcript provides an independent geometric validation of the CellDot correction. Corrected transcripts were matched back to the platform transcript table by their coordinates and compared across fates using three measures: nuclear overlap, distance to the nucleus, and, for reassigned transcripts, whether they were located on the side of the original cell facing the destination cell. At the tissue level, we quantified the expression similarity between adjacent cells from different compartments and measured the tumor-marker signal in non-tumor cells as a function of the distance to the nearest tumor cell. We additionally performed a model-free directional analysis independent of the CellDot output by testing whether misplaced marker transcripts preferentially faced the nearest cell of the corresponding marker-expressing compartment, using correctly localized marker transcripts as a control.

### Cell-cell communication analysis

Ligand-receptor interactions were inferred with three tools, including CellPhoneDB (v5, database v5.0.0)[34] with its statistical method (1,000 permutations, expression threshold 0.1, *P* < 0.05), CellChat (v2.2.0)[35] with its default permutation test and a minimum of 10 cells per type, and the rank-aggregate consensus of LIANA+[36] (v1.7.3, expression proportion 0.1, significance at specificity rank ≤0.05). Each tool ran with identical settings on three inputs, namely the raw data, the corrected data and the dissociated single-cell reference restricted to the same panel and cell types, all as log-normalized counts. Interactions were compared with the reference at two levels, including the pair level, which asks only whether a ligand-receptor pair is detected, and the edge level, which additionally requires the sending and receiving cell types to match. Precision is the fraction of detected interactions also found in the reference, and recall the fraction of reference interactions recovered.

### Analyses of the whole-transcriptome section

#### Gene groups and cell-type specificity

Genes were grouped according to whether they were included in the 5,101-gene Xenium 5K panel of a cervical cancer Xenium Prime dataset, which is the 5,001-gene Xenium Prime 5K panel with a 100-gene add-on, based on the gene-panel file released with that dataset. For each gene, we computed the cell-type specificity score *τ* of Yanai et al.[40] based on the single-cell reference. Let *P*[*t, g*] denote the reference expression profile of gene *g*, obtained by pooling counts within each of the *T* = 17 types and normalizing each type to counts per ten thousand. Cell-type specificity[40] was then defined as *τ*_*g*_ = ∑_*t*_(1−*P*[*t, g*]/ max_*t*_′ *P*[*t*^′^, *g*])/(*T* −1), where *τ*_*g*_ = 0 indicates uniform expression across cell types and *τ*_*g*_ = 1 indicates expression restricted to a single cell type. Using the molecule-level fate table, we calculated the fraction of transcripts reassigned for each gene and summarized its relationship with *τ* by the median reassignment rate across eleven equal-width *τ* bins, restricting the analysis to genes with more than 200 detected transcripts. For the comparison of *CD48*, we calculated the mean log-normalized expression of each cell type in the reference, the raw and the corrected data, using up to 12,000 cells per type.

#### Spatial niche analysis

Niches around B cells were identified with NicheScope[41] on the raw and on the corrected layer independently. Each cell’s neighborhood is summarized as a Gaussian-weighted cell-type composition (*σ* = 20 µm), and NicheScope couples this composition with the cell’s own expression (3,000 highly variable genes, three components), returning per niche a gene program and a neighbor cell-type profile together with their canonical correlation. In each layer, the component whose strongest neighbor weight is assigned to a T-cell type was taken as the immune niche around B cells. The composition panel shows the cell types with the largest neighbor weights, and the program panel shows the genes with the highest loadings. Gene Ontology biological-process enrichment was performed on the 120 top program genes of each fit using Enrichr[50] and the GO Biological Process 2023 library, with immune-related terms ranked by adjusted *P* value. In the molecule-level analysis, a corrected transcript was counted when CellDot reassigned it from a cell of another type to a cell type in which the gene is expressed.

## Data availability

### Spatial datasets

Four Xenium datasets were included in this study, namely the human breast cancer (167,780 cells, 313-gene panel; https://www.10xgenomics.com/products/xenium-in-situ/preview-dataset-human-breast), the human colorectal cancer (340,837 cells, 422-gene panel, sample P2; https://www.10xgenomics.com/platforms/visium/product-family/dataset-human-crc), whose matched Flex, Visium and Visium HD data from the same specimen serve as external verification, the human lung cancer (278,328 cells, 5,001-gene Xenium Prime 5K panel; https://www.10xgenomics.com/cn/datasets/xenium-human-lung-cancer-post-xenium-technote), and the human cervical cancer profiled with the Atera whole transcriptome assay (717,576 cells, 18,028-gene panel; https://www.10xgenomics.com/datasets/atera-wta-ffpe-human-cervical-cancer).

### Single-cell references

Each spatial dataset is paired with a labeled single-cell reference. The breast section uses the matched single-cell FFPE dataset released with the same study[4]; the colorectal section uses the Flex dataset of the same specimen from the family stated above; the lung section uses the public LuCA lung cancer atlas[51] (https://cellxgene.cziscience.com/collections/edb893ee-4066-4128-9aec-5eb2b03f8287). The cervical section uses the paired cervical scFFPE dataset (https://www.10xgenomics.com/datasets/17k-human-cervical-cancer-scFFPE), which is released without labels and was annotated de novo by Leiden clustering followed by canonical-marker scoring per cluster; because dissociation depletes neutrophils, a neutrophil profile from the LuCA atlas above was added as one further reference type. CellDot operates on the genes shared between each panel and its corresponding reference. Dataset summaries and per-dataset preprocessing are provided in Supplementary Table 2 and Supplementary Note 5.

### Processed data

The CellDot outputs on the four sections (the corrected cell-by-gene matrices, the fate of every molecule and the references used), together with the data behind every panel of the main figures, are deposited on Zenodo (https://doi.org/10.5281/zenodo.22489061).

## Code availability

CellDot and the tutorials for reproducing the analyses are available at https://github.com/YangLabHKUST/CellDot. The code that reproduces every panel of the main figures, one script per panel with the plotting style of the paper and the scripts of the analyses behind each panel, is archived with its data on Zenodo (https://doi.org/10.5281/zenodo.22489061). An interactive viewer of the CellDot results on the four datasets in this paper is available at https://viewer.celldot.online/.

## Acknowledgements

This work was partly supported by the Young Student Basic Research Program under the National Natural Science Foundation of China (125B2027); the Innovation and Technology Commission (ITC-SKLNSD26SC01); Hong Kong Research Grants Council Grants, AoE/E-601/24-N, C6040-24G, 16308120, 16307221, 16307322, 16302823, 16309424, and 16308925; The Hong Kong University of Science and Technology Startup Grants R9405 and Z0428 from the Big Data Institute. The computation tasks for this work were performed using the X-GPU cluster supported by the Research Grants Council Collaborative Research Fund Grant C6021-19EF and HKUST SuperPOD. J.X. was supported by National Natural Science Foundation of China (Grant No. 12401384); Guangdong Natural Science Foundation General Project (Grant No. 2025A1515011603); and Sun Yat-sen University Startup Grant. A.R.W. was supported by the National Natural Science Foundation of China (T2422018). The funders had no role in study design, data collection and analysis, the decision to publish, or preparation of the manuscript.

## Author contributions

C.Y., Y.L. and Y.C. conceived the study. C.Y. and J.X. supervised the project. Y.C. designed, implemented and validated CellDot. J.X. and Y.L. contributed to the experimental design. Y.Z., S.H., Z.C. and B.Y. assisted with the validation of CellDot. F.Z., A.R.W., H.C. and J.W. helped with analyzing the results. Y.C., Y.L. and C.Y. wrote the manuscript with input from all the authors.

